# Coordinated multivoxel coding beyond univariate effects is not likely to be observable in fMRI data

**DOI:** 10.1101/2021.06.13.448229

**Authors:** Mansooreh Pakravan, Ali Ghazizadeh

## Abstract

Simultaneous recording of activity across brain regions can contain additional information compared to regional recordings done in isolation. In particular, multivariate pattern analysis (MVPA) across voxels has been interpreted as evidence for distributed coding of cognitive or sensorimotor processes beyond what can be gleaned from a collection of univariate responses (UVR) using functional magnetic resonance imaging (fMRI). Here, we argue that regardless of patterns revealed, conventional MVPA is merely a decoding tool with increased sensitivity arising from considering a large number of ‘weak classifiers’ (i.e. single voxels) in higher dimensions. We propose instead that ‘real’ multivoxel coding should result in changes in higher-order statistics across voxels between conditions such as second-order multivariate responses (sMVR). Surprisingly, analysis of conditions with robust multivariate responses (MVR) revealed by MVPA failed to show significant sMVR in two species (humans and macaques). Further analysis showed that while both MVR and sMVR can be readily observed in the spiking activity of neuronal populations, the slow and nonlinear hemodynamic coupling and low spatial resolution of fMRI activations make the observation of higher-order statistics between voxels highly unlikely. These results reveal inherent limitations of fMRI signals for studying coordinated coding across voxels. Together, these findings suggest that care should be taken in interpreting significant MVPA results as representing anything beyond a collection of univariate effects.

## 1. Introduction

Multivariate pattern analysis (MVPA) has received much attention as a popular decoding tool for fMRI studies [1]. In classical fMRI analysis methods, individual voxel responses to task events are analyzed independent of other voxels (univariate analysis or UVA) using generalized regression [2, 3]. However, in a typical MVPA study, the goal is to investigate whether it is possible to decode task conditions or states from fMRI signals of a collection of voxels in a given brain area [4, 5, 6, 7]. Local MVPA aims to assess the task-related population code by analyzing the fMRI signals extracted from a local neighborhood (searchlight) surrounding each voxel [6, 8, 9].

A key reason for the wide adoption of MVPA is its success in identifying multivoxel neural representations of task conditions[10]. Indeed, by combining information across voxels, MVPA increases the dimensionality for the optimal criterion thereby improving the decoding performance with higher sensitivity and specificity [11, 12]. In some cases regions that do not show significant UVR are found to have significant task-related information when MVPA is used. In such a scheme, each voxel may have a weak bias across task conditions due to weak neural activation or biased sampling of neurons with different tuning properties concerning task conditions [13]. Such weak biases which may be missed by UVA as being sub-threshold can nevertheless be pooled across voxels for successful decoding in MVPA in higher dimensions determined by the number of voxels considered[14, 15, 16].

However, beyond improved decoding performance compared to UVA, there is an ambiguity as to how much of the observed multivariate patterns differences across conditions is the trivial reflection of univariate response differences. The concern for univariate response removal is valid because in many cases it is not clear whether multidimensional representations in the brain can be explained by a collection of ‘simple’ unidimensional activations or not [17]. To address this issue, various approaches for removal of UVR components have been suggested including removal of mean response across voxels in each condition [18, 19, 20, 21] or removal of common pattern across conditions [22].

Here we argue that none of these approaches can help determine whether there is joint multivoxel coding beyond univariate effects. This is because MVPA cannot capture the information contained in voxels beyond changes in the mean across conditions. Thus there are no fundamental differences between the precise nature of decoded information using single-voxel activations in UVA and multi-voxels activations in MVPA (both relying on changes in mean responses across conditions). To find joint multivoxel coding beyond univariate effects one should look for significant changes in higher-order statistics of voxel activations. To address this issue, fMRI datasets in both human and non-human primates with significant multivoxel responses using MVPA were analyzed for the presence of such higher-order statistics (here second order). Surprisingly, results showed a lack of significant modulation of second-order statistics by task conditions despite significant univariate and multivariate coding based on mean activity and despite the fact that such second-order effects were readily observable in population neuronal spike data consistent with previous reports [23, 24, 25]. Further analysis suggested that the difficulty of detecting higher-order effects might be inherent to fMRI and thus unavoidable due to nonlinear hemodynamic coupling and low spatial resolutions of fMRI signals.

## 2. Results

## 3. Univoxel versus multivoxel responses

Better classification performance using MVPA can be a trivial consequence of the fact that classification is always a monotonically non-decreasing function of the number of voxels in a searchlight (Fig 1A). As can be seen increasing the number of voxels results in larger Crossnobis distance [18] across two conditions and thereby improving classification accuracy for any classifier such as linear discriminant analysis (LDA) [21] (see section 8) (Fig 1B). It can be argued that almost all previously reported MVPA results could arise from a mere increase in sensitivity which is expected when combining a large number of ‘weak classifiers’(i.e. single voxels) within searchlights and solving the classification in a higher-dimensional space [14, 15, 26]. While such results maybe used as evidence for the spatial spread of coding beyond a single voxel, they hardly address whether there is coordinated activity between voxels that are different across conditions. Note that the spatial spread of coding are also often readily observed using simple univariate GLMs unless voxels in the searchlight show mostly subthreshold activations that can only reach significance when MVPA is used (again via increasing sensitivity by using a large number of ‘weak classifiers’)

**Figure 1:**
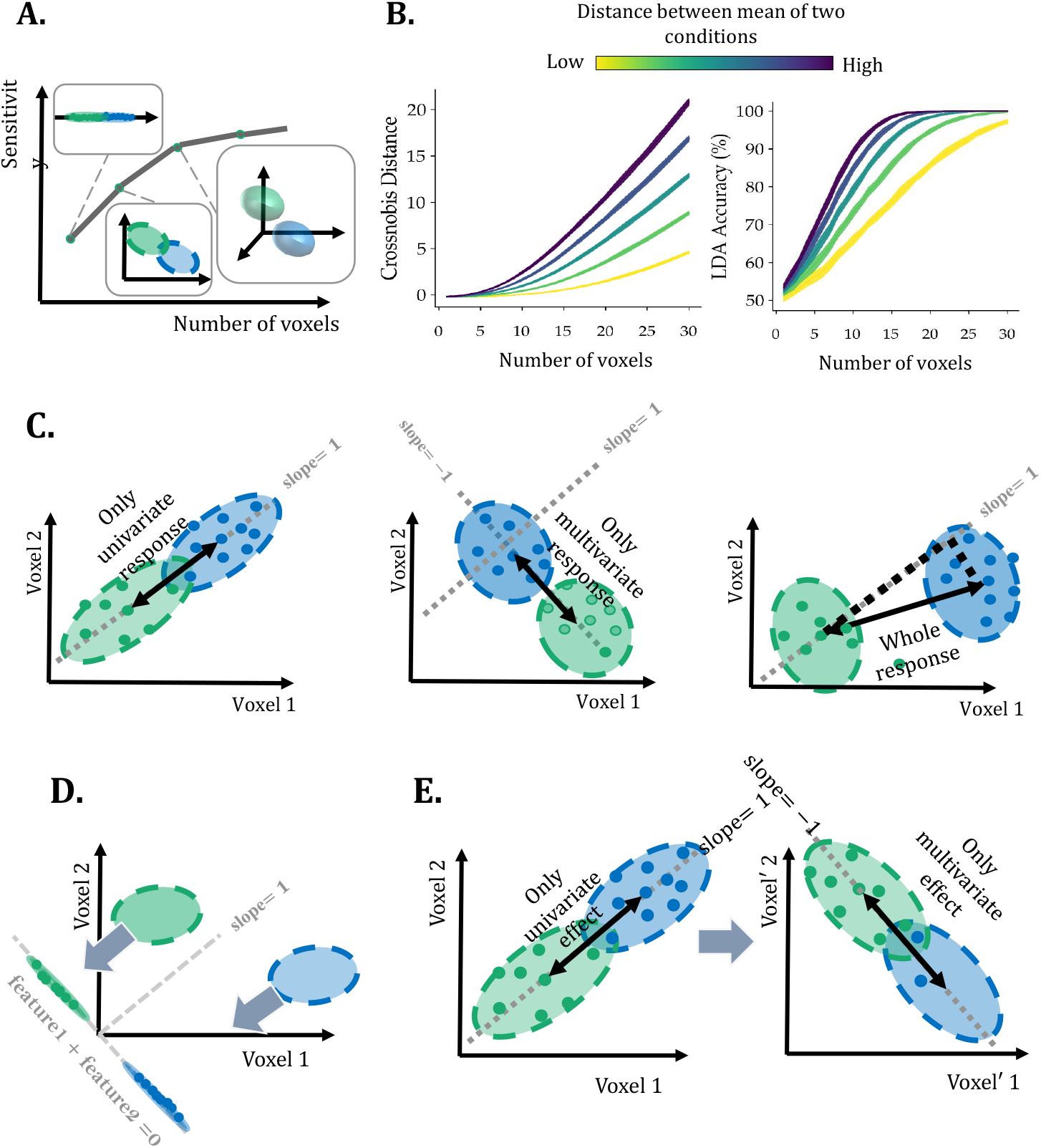
UVR and MVR components and sensitivity to the number of voxels. (A) MVPA accuracy in condition discrimination (sensitivity) increases by aggregating information across larger number of voxels (B) As the number of voxels increases, the Crossnobis distance (see section 8) among conditions increases and so is classification accuracy by linear discriminant analysis (LDA) for any given structure of noise. (C) The classical view of UVR and MVR components when doing MVPA: UVR is the component along the unity line of equal sensitivity across voxels (left) and MVR is the component perpendicular to this line (middle). Voxels can show a mixture of UVR and MVR in this scheme (right). (D) The conventional approach in removing MVR from UVR by projecting the data onto the hyper-diagonal of voxel space for each condition separately. (E) Equivalence of classical definition of UVR and MVR by affine transformation and in terms of informational content for discriminating two conditions.

As a result in many studies, it has been a challenge to determine to what degree the results of MVPA are simply a byproduct of UVR as opposed to representing a real MVR code [19, 1]. Several studies have assumed that in general if voxels show similar activation across conditions (e.g. increased or decreased activation in all voxels between two conditions), MVPA is simply a byproduct of multiple UVRs [1, 19] and on the other hand, when there is differential activity across voxels across task conditions, the activation pattern is assumed to represent a real MVR code. Fig 1C shows a graphical representation of UVR and MVR according to the aforementioned interpretation for two voxels across two conditions. MVPA activations are interpreted as UVR when activation changes are largely similar for both voxels across the two conditions (i.e. activation falling on the unity line which represents equal sensitivity, UVR axis). In this scenario, MVR is a component that is added to the UVR in the direction perpendicular to the UVR axis. In reality, the actual activation of our two example voxels can show a mixture of both UVR and MVR (Fig 1C right). A conventional method employed to remove UVR from MVR is then to remove the part of the response that has the same sign and amplitude across all voxels [1, 18] (Fig 1D). This procedure is separately performed for each condition. In the geometric sense, MVR is extracted by shifting all samples in each condition such that the sum of voxel values for each sample becomes zero [1].

Unfortunately, such interpretation of MVPA can be erroneous for at least two reasons: 1) Current UVR interpretation implicitly assumes equal sensitivity to conditions across voxels. However, it is well-known that a given voxel’s sensitivity to task conditions can be affected by a variety of factors not related to neural activity per se, including imaging sequence used, RF coil function and positioning, subject motion as well as other artifacts and dropouts common in an imaging session. In the interpretation proffered in Fig 1 such differential sensitivity across voxels can be taken as evidence for MVR in the absence of a meaningful neural correlate. 2) More importantly from a computational standpoint and as far as informational content or decoding is concerned, conditions with pure UVR and pure MVR (current formulations as shown in Fig 1C) are not qualitatively different. In our two voxel examples, pure UVR can be transformed to pure MVR by an affine transformation of voxel activities (Figs. 1E). Importantly and as expected such linear transformations do not change the informational content or decoding accuracy regardless of which name (UVR or MVR) one chooses for describing the observed pattern.

On the other hand, we argue that a true coordinated and joint multi-voxel code which posits functional cross-voxel interactions across conditions entails changes in higher-order cross-voxel statistics. Such cross-voxel interaction entangles response fluctuations across voxels within trials. This alternative formulation is consistent with a more precise definition which suggests that a ‘true’ population code should be represented in coordinated activity between members within a given trial and not just their mean across trials [27].

## 4. MVR vs sMVR

In theory, arbitrary interactions between any number of voxels can be considered resulting in higher-order interaction (between 2 to ‘N’ voxels) however correct estimate of such higher-order terms suffers from the curse of dimensionality and requires increasingly larger amounts of data. The least amount of complexity that still goes beyond mean responses is to consider pairwise covariance of voxel activations (sMVR).

Thus, for a condition indicator *x* and two voxels *v*_1_ and *v*_2_, formulating the decoding as a simple additive model with a pairwise interaction term can be written as:

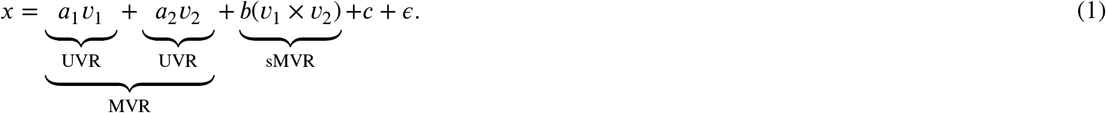

where *c* and *є* are constant and residuals, respectively.

Note that the classical MVPA follows basically the same model but without the interaction term *d*(*v*_1_ × *v*_2_). Each term *f*_1_ or *f*_2_ in Eq. (1) can be considered UVR while the sum (*av*_1_+*bv*_2_) can be considered MVR which is examined by MVPA and nevertheless has the same nature of information as the UVR terms. On the other hand, the term *d*(*v*_1_ × *v*_2_) which represents the second-order interaction effect can be considered as the simplest form in which coordinated activity across voxels may be used for discrimination of two conditions. This is the term that we refer to as second-order MVR (sMVR). The expected value of this second-order interaction is the covariance matrix of voxels in the searchlight provided that the activations are all mean subtracted. Significant changes in the covariance matrix can reveal additional information in the voxels about the different conditions beyond their mean activation. Thus, in sMVR, the discrimination of conditions depends on how well the covariance structure of the data varies across conditions.

There are different ways to measure MVR including examining the accuracy of classifications [5] or the continuous distance measures such [28]. In this study, squared cross-validated Mahalanobis distance that is also known as Crossnobis distance is used to extract the strength of MVR [28, 18] (Fig S1). Notably, while classification accuracy is a discrete measure and saturates at 100%, the Crossnobis distance is a continuous and non-saturating measure and thus allows for better differentiation of cases with very high classification accuracies [1, 28]. To extract sMVR, Geodesic distance that is a geometry-aware approach to compare covariance matrices is applied [29] (Fig S2) (see section 8 for more details).

Fig 2A illustrates a simplified view of MVR and sMVR for two voxels in four different scenarios using simulated data. If the covariance matrices of two conditions are different (scenarios 2 and 3), then there is sMVR and if the mean activities are different, then there is MVR (scenarios 1 and 2). Fig 2B shows one sample in two contexts (say *A* and *B*) across matrix of voxels with 4 sub-patches (each consisting of 9 voxels) coding the conditions *A* and *B* with MVR only (subpatch #1), sMVR only (subpatch #3), MVR+sMVR (subpatch #2) or no coding (subpatch #4) same as the scenarios considered in Fig 2A. As can be seen in this proof of concept simulation, Crossnobis and Geodesic distances showed differential sensitivity to subpatches with MVR and sMVR coding, respectively (Fig 2C).

**Figure 2:**
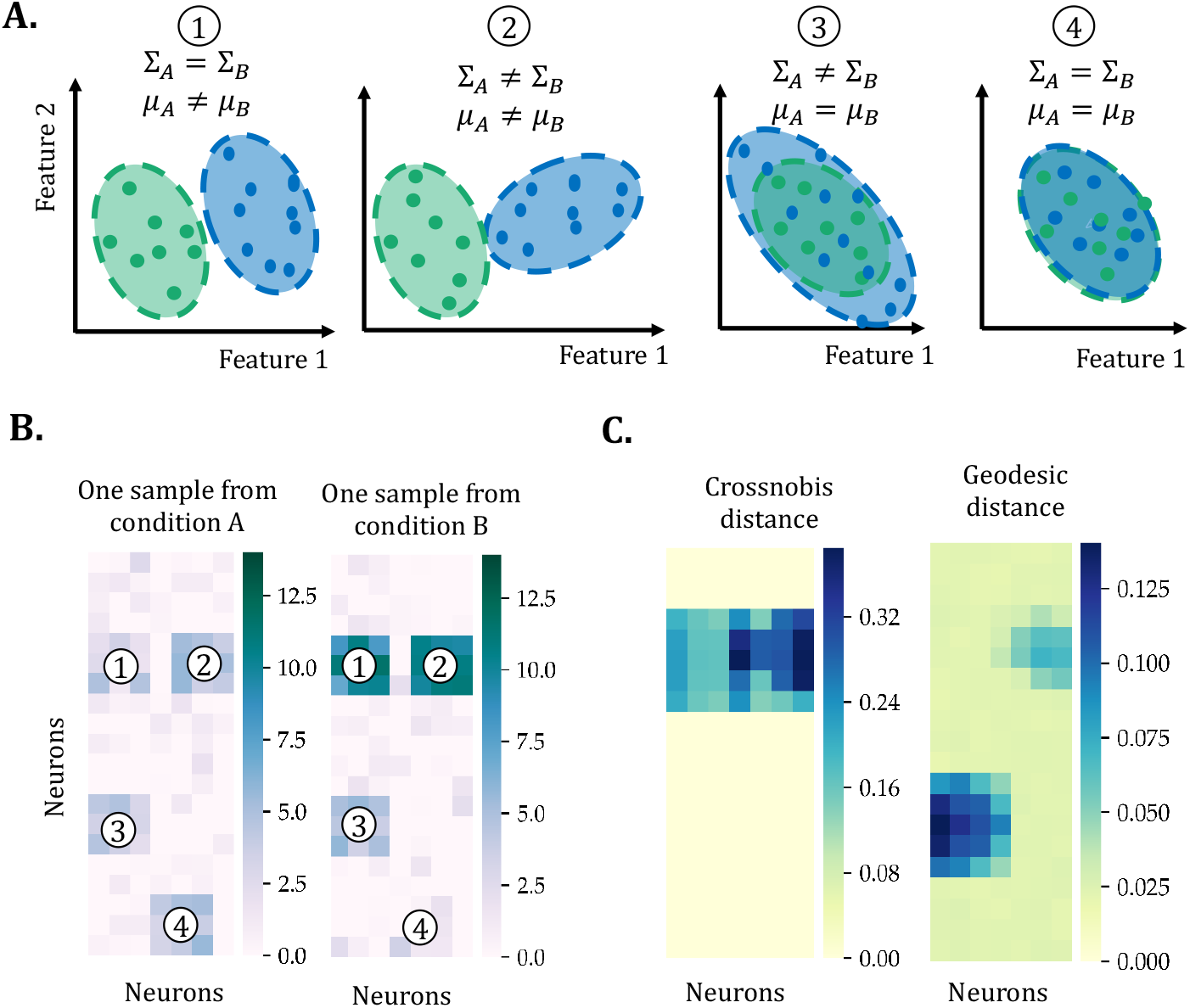
Crossnobis and Geodesic distance are differentially sensitive to changes in population mean and covariance matrix, respectively. (A) Double dissociation of MVR and sMVR responses: Scenario 1, has MVR but no sMVR, scenario 2 has both MVR and sMVR, scenario 3 has sMVR but no MVR, and scenario 4 has neither MVR nor sMVR. (B) A sample matrix of voxels in conditions A and B with patches corresponding to the four different scenarios in A. (C) Crossnobis distance correctly detects scenarios 1 and 2 which have MVR effects regardless of sMVR. On the other hand, Geodesic distance correctly detects scenarios 2 and 3 with sMVR regardless of MVR. (for this simulation the number of whole voxels was 140 with 500 samples.

## 5. No sMVR in fMRI data for conditions with significant MVR in two species

Previously published fMRI datasets across human and non-human primates (macaques) were analyzed in this study [30, 31, 16]. In the macaque dataset, two adult rhesus monkeys (D and U) participated in the recording sessions. In one experiment, each monkey had 12 face-patch localizer runs with each run consisting of trials in which the monkey passively viewed centrally presented images of conspecific monkey faces or ordinary objects. In another experiment, computer-generated fractals were presented in the left or right visual hemifields in a pseudorandom order. In the human fMRI dataset, 17 subjects were recruited each with two face/scene localizer runs [16].

Fig 3A illustrates MVR results (Crossnobis distance) for two monkeys across two different task conditions (face vs object or left vs right hemifield fractal presentation) using a searchlight with 15 voxels. The face vs object Crossnobis distance nicely marked the well-known face patches in the ventral stream including areas PL, ML/MF, and AL/AF [32]. The left vs right hemifield conditions mainly marked posterior visual areas including V1-V4 and TEO which are known to have strong laterality in their responses [33]. Notably, both results are in good agreement with previous general linear model (GLM) analysis [30] of face vs object and left vs right using the same data set (Fig S3).

**Figure 3:**
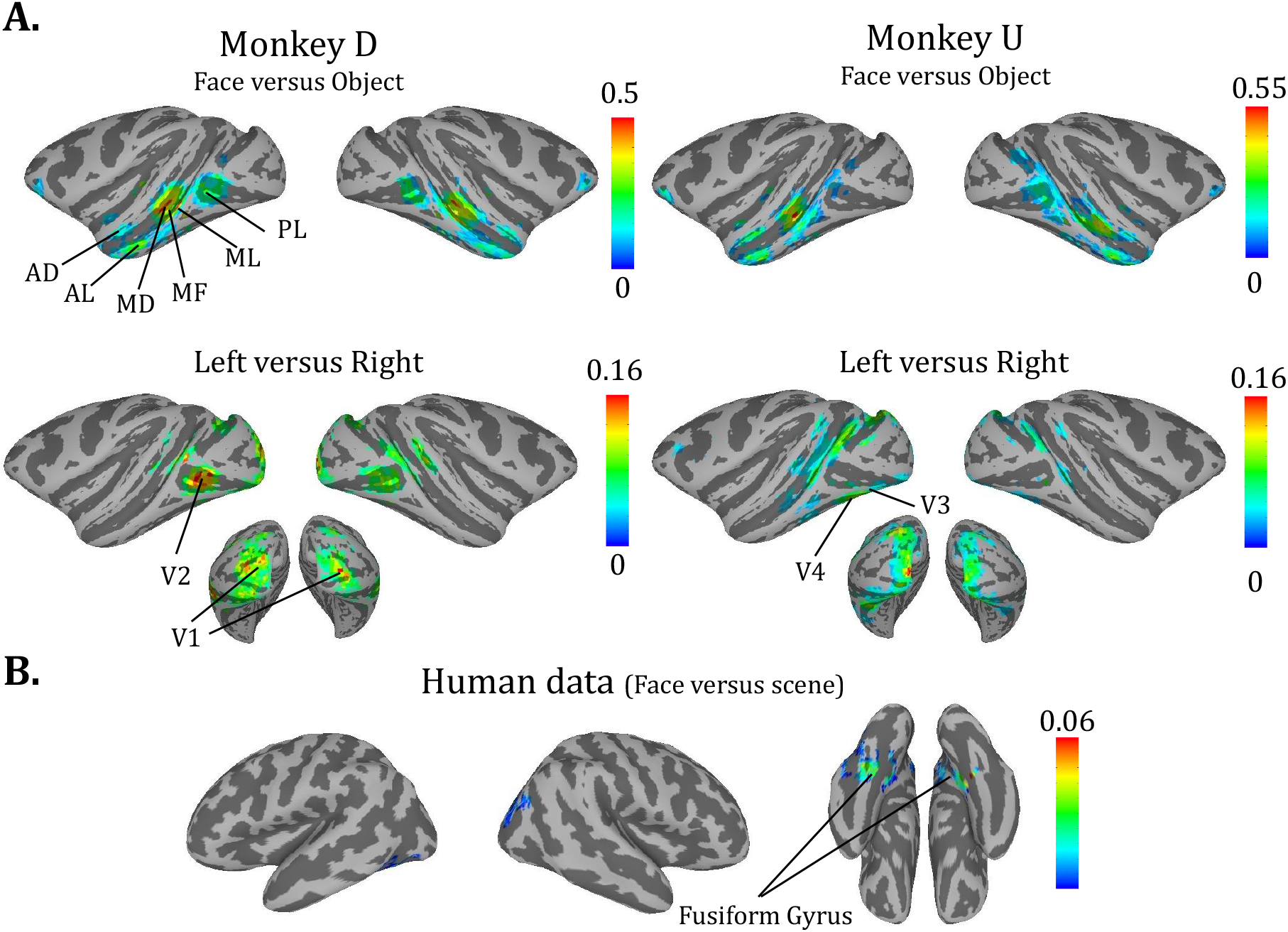
MVR for face/object and left/right hemifield across cortical areas (A) Crossnobis distance (with 15 voxels in each searchlight) for two monkeys for face/object and left/right discrimination tasks. The highlighted regions show significant cluster of voxels (*P*_*value*_ *<* 0.01, minimum cluster size 30) including areas PL, ML/MF and AL/AF for face/object and left/right in V1-4 and TEO areas for left/right contrasts. (B) The average of Crossnobis distance of 17 subjects for face/scene discrimination task. The highlighted regions are significant cluster of voxels (*P*_*value*_ *<* 0.01, minimum cluster size 30) situated in Fusiform Gyrus.

Fig 3B shows the group analysis results across 17 human subjects for face vs object contrast which shows robust MVR in the fusiform area again in agreement with previous literature using univariate approaches ([34]). In both human and monkey datasets the MVR results tended to show wider activation compared to univariate analysis as expected due to the higher sensitivity of the multivariate analysis (Fig 3, Fig S3 and Fig S4).

Fig 4 shows the Geodesic distance of different brain regions for two monkeys with the same two task conditions face/object and left/right considered previously (Fig 4A) and the average Geodesic distance of 17 subjects in face/object conditions (Fig 4B). Notably, we did not find any cluster of voxels with significant changes in the second-order statistics across conditions in either species. To make sure that meaningful clusters with sub-threshold activations were not missed, voxels with the top 5% sMVR values for monkeys data and 20% sMVR values for human data were marked. Examination of activated regions showed a dispersed pattern across the cortex for the two monkeys Fig 4A and for group-level human data Fig 4A-B. Furthermore, the fact that the areas with larger sMVR values did not show a systematic difference between two tasks (face/object and left/right) in the two monkeys further casts doubt on their functional significance (4A). In addition, in comparison to highly consistent effects of MVR (Crossnobis distance) across subjects, the areas with high sMVR was much less consistent across human subjects (Fig S5).

**Figure 4:**
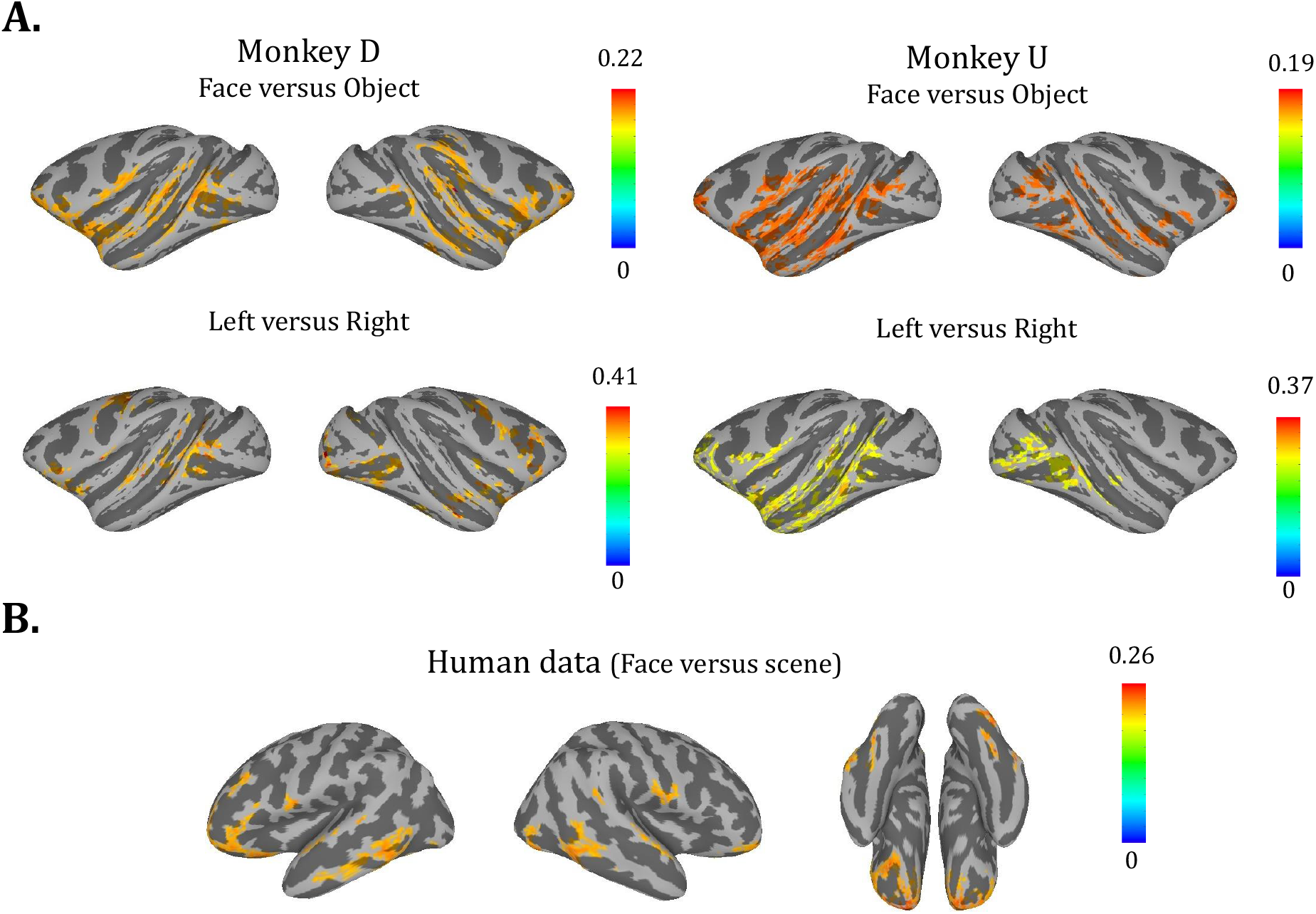
sMVR for face/object and left/right hemifield across cortical areas (A) Geodesic distance (with 15 voxels in each searchlight) for two monkeys for face/object and left/right discrimination tasks. (B) The average of Geodesic distance for 17 subjects for face/scene discrimination task. The highlighted regions show top 5% and 20% regions with highest sMVR (minimum cluster size=30) in monkeys and human dataset, respectively. Note that none of these clusters were significant (*P*_*value*_ *>* 0.01).

To further examine whether regions with significant MVR coding of face/object or left/right showed any significant second-order changes (i.e. sMVR), distribution of pairwise voxel values across two conditions were plotted in two face patches (MD and AL) and two regions with strong laterality coding (Fig 5). Fig 5A shows this pairwise distribution in both monkeys in areas MD and AL. As expected given significant Crossnobis values, the mean of the two distributions was different in these regions in both monkeys (shift in the center of covariance ellipses). However, there was minimal change in the shape or orientation of the covariance ellipse. Fig 5B shows the same analysis for areas V1 and V4 which showed significant MVR for left/right conditions but no significant sMVR. Fig S6 visualizes the changes in the covariance of voxels across left versus right conditions by removing the mean of each condition showing the largely similar covariance matrices across conditions. The distribution of pairwise voxel values across two conditions in the human dataset is also presented in Fig S7.

**Figure 5:**
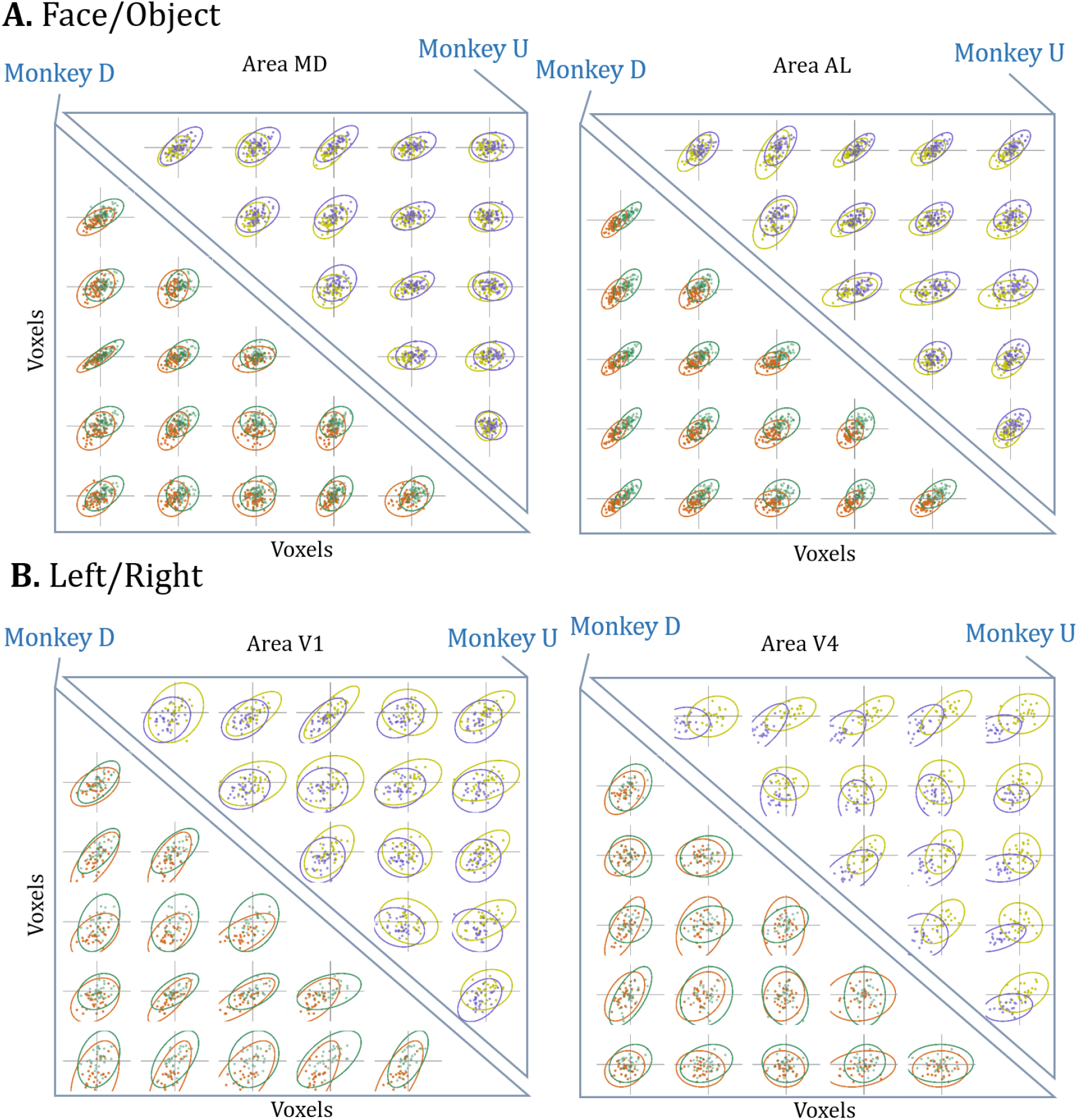
Visualization of changes in mean vs covariance of voxels across conditions. (A) Joint-distribution of pairwise voxel activations in monkey D (lower triangle) and monkey D (upper triangle) in face vs object blocks for two different searchlights in area MD (left) and area AL (right) with significant Crossnobis distance and non-significant Geodesic distance. (B) same format as A but for left vs right blocks in area V1 (left) and area V4 (right). Plots in panels A and B are represented in Fig S6 without mean differences between conditions.

Thus the examined contrasts failed to show evidence for modulation of coordinated voxel activity by task conditions in either species even in cases where both conditions evoked significant MVR across the cerebral cortex.

## 6. Can sMVR be observed when there is second-order population response at the neural level?

We previously showed that Geodesic distance can successfully detect sMVR in simulated data (Fig 2B-C). However, our analysis did not find evidence for sMVR in the examined fMRI datasets. To make sure that Geodesic distance can detect second-order effects in real neuronal data, the same approach was applied to recorded neuronal populations in primary and secondary visual cortices (area V2) in response to gratings with various orientations. Here, results showed significant coding in both the mean responses (Crossnobis distance) as well as in the second-order effects (Geodesic distance) in the neural population (Fig 6). Interestingly, both Crossnobis and Geodesic distances varied in a similar fashion as a function of the angle between the two gratings shown reaching their maximum at the 90-degree difference. This fact can be verified by looking at the difference in the mean response and the covariance visualized for an example set of neurons (Fig S8).

**Figure 6:**
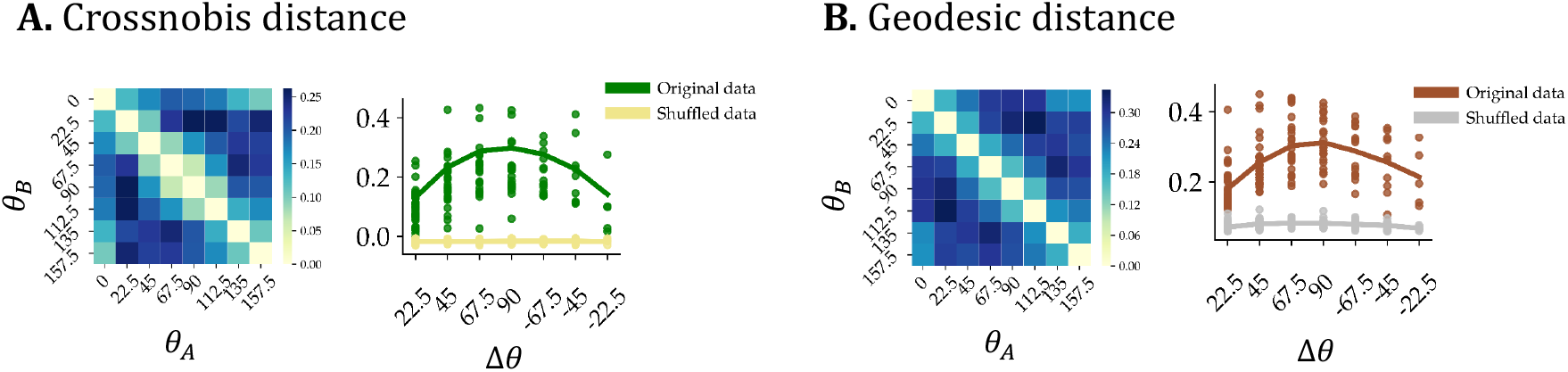
Detected UVR and MVR in recorded neuronal populations in primary and secondary visual cortices. (A) The left two plots: average of Crossnobis distance between pairs of *θ*_*A*_ and *θ*_*B*_ in visual area V2. (*θ*_*A*_ and *θ*_*B*_ are grating degree of stimulus for conditions *A* and *B*, respectively and the the plots are the average of 5 recorded sessions). The right two plots: Crossnobis distance versus Δ *θ* = *θ*_*A*_ − *θ*_*B*_ for all sessions. (B) same format as A but for Geodesic distance. Note that the maximum absolute value of Δ *θ* is 90. The results of original data are significantly higher than shuffled data for all (*P*_*value*_ *<* 0.01).

However, the presence of sMVR in the neuronal population does not guaranty its detectability in the associated fMRI signals. Indeed a reasonable expectation is that due to temporal averaging and distortions caused by nonlinear hemodynamic coupling and the spatial integration of neural responses across large populations with possibly heterogeneous responses across each condition within each voxel, the effects of first-order and second-order modulations (i.e. MVR and sMVR) could be smeared to a large extent. To examine such effects, predicted fMRI activations arising from V2 populations were simulated using forward hemodynamic model with spatial smoothing (Fig 7A) (see section 8 for more details).

**Figure 7:**
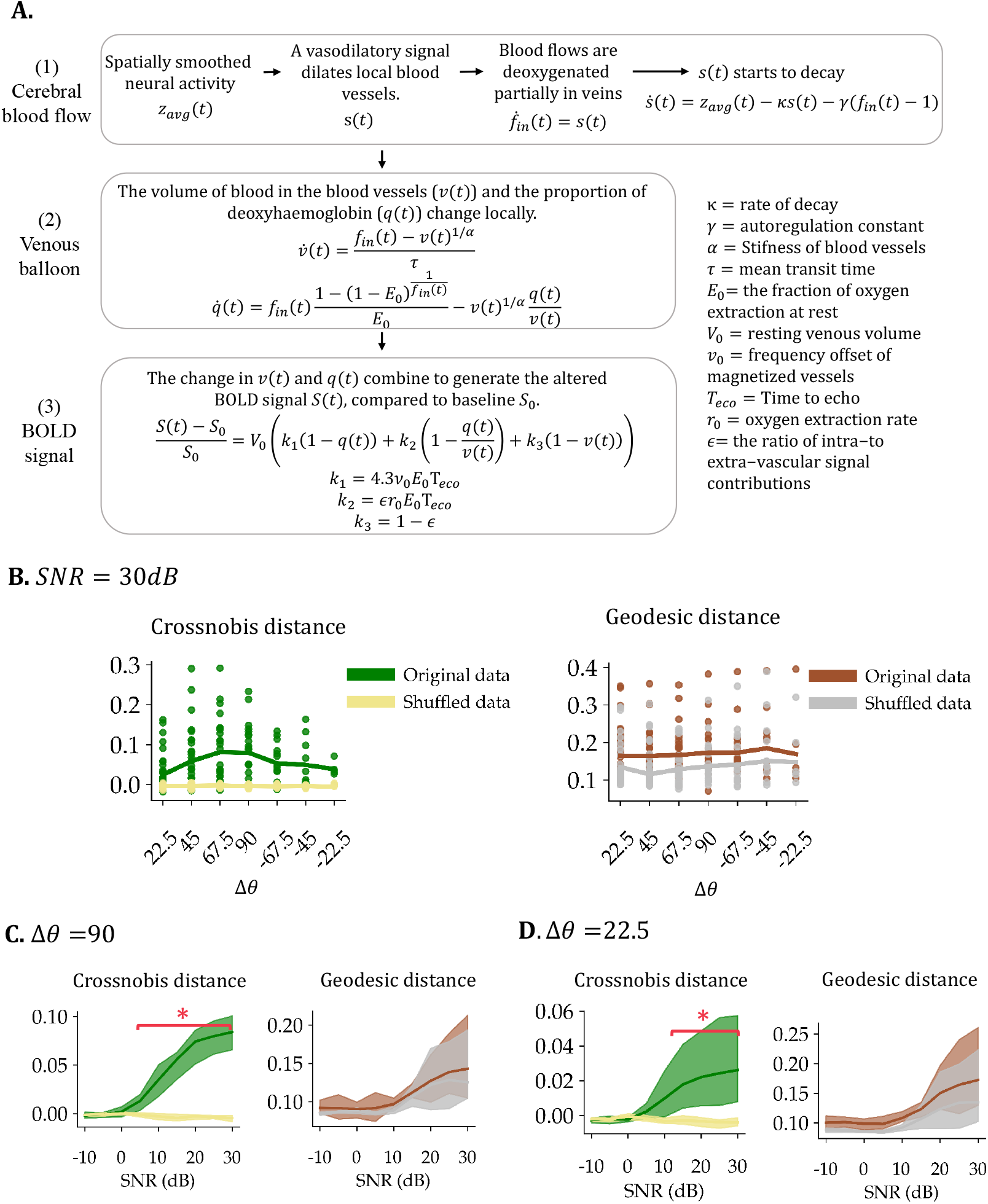
Simulated BOLD signal from neural activity using forward hemodynamic model and diminished spatial resolution (A) A tripartite hemodynamic forward model used to transform neural activity *z*(*t*) to the BOLD signal *S*(*t*). First, neural activity *z*(*t*) triggers a vasolidatory signal *s*(*t*) which leads blood flow to increase. Second, the flow leads a change in both blood volume *v*(*t*) and deoxyhaemoglobin *q*(*t*) and third both *v*(*t*) and *q*(*t*) combine non-linearly to generate BOLD signal *S*(*t*). (B-D) Effect of hemodynamic coupling and spatial averaging within voxels in BOLD images on detectability of MVR and sMVR (B) Crossnobis and Geodesic distances of simulated BOLD signal using real neural data in V2 with 30dB SNR. (C) Crossnobis and Geodesic distances for noisy simulated BOLD signal given real neural data in V2 as a function of increasing SNR for *Δ θ*= 90. (D) same format as C but for *Δ θ*= 22.5. The SNRs with statistically significant differences are marked with a red line and asterisk (*P*_*value*_ *<* 10−3). Geodesic distances in none of the scenarios were non significant.

Fig 7B presents Crossnobis and Geodesic distance of simulated BOLD signal assuming a high signal-to-noise (SNR: 30 *dB*) with additive noise added to the actual V2 neural data. As it can be seen, information transfer from spikes to BOLD is concomitant with a significant reduction of discriminability in both MVR and sMVR but with a more deleterious effect on the second-order effects. In this case only MVR were significantly detectable in the simulated BOLD data with dependence on *Δθ* reaching a maximum at *Δθ* = 90 (*P*_*value*_ *<* 0.01). Notably, neither region showed significant sMVR despite robust second-order effects in the neural spiking data (Fig S9). As expected the detectability of both MVR and sMVR was an increasing function of SNR (Figs. 7C-D). However, while MVR could be detected at moderate SNRs (10-20 *dB*) in both regions, sMVR was hard to detect even at very high SNRs (30 *dB*). Examination of joint-distribution of pairwise voxel activations using this simulated data also visually confirms this lack of sMVR even at a high SNR and even for large differences in conditions (*Δθ* = 90) (Fig S9). The difficulty in finding significant effects especially for low SNRs readily explains the lack of sMVR modulation in real fMRI data (Fig 4) which have typically low SNRs *<* 10*dB* (Fig S10) despite significant MVR coding (Fig 3). Note that the macaque fMRI data presented were obtained with MION which has slower dynamics than BOLD but slightly better spatial specificity. Nevertheless, the intuitions from the hemodynamic model used should more or less be applicable to the MION signal as well [35].

It is worth mentioning that the synthesized BOLD signal depends on the model’s parameters, which generally have some useful interpretations. Changing various hemodynamic parameters had not any significant effect on the Geodesic distance of synthesized fMRI data.

## 7. Discussion

MVPA has found widespread applications in fMRI data analysis but it has critical limitations for parsing multivariate coding arising from correlated voxel activations beyond what can be gleaned from a collection of activations in individual voxels. Here, we first presented a methodology for double dissociation of first and second-order effects of conditions on voxel activations (Crossnobis and Geodesic distances for quantifying MVR vs sMVR respectively). Surprisingly, analysis of conditions eliciting robust UVR and MVR coding in the brain (face/object and left/right hemifield) failed to show significant sMVR using whole-brain fMRI in two species (humans and macaques). Further analysis showed that while both MVR and sMVR can be readily present in the spiking activity of neural population, the slow and nonlinear hemodynamic coupling and low spatial resolution of fMRI activation can obfuscate the second-order effects to a large degree.

In MVPA studies, the meaning of “removal of UVR” and what remains after this removal is not exactly clear [19, 17, 36]. Here, we argued that conventional MVPA which is based on the comparison of multivoxel means across conditions is inherently incapable of finding ‘true’ joint multivoxel coding that should be reflected in changes in the coordinated activity of voxels across conditions. Some of the previous suggestions for separating UVR and MVR are only parsing uniform and non-uniform mean response differences to conditions [1]. Here, we suggest that any decoding done with more than one voxel should be considered MVR and that there is no inherent difference in decoding of conditions for mean responses falling on the unity line or orthogonal to it in the high dimensional voxel space 1C). While conventional MVPA is affected by noise correlations across voxels [16], it does not explicitly track changes in noise correlation structure across conditions. Instead, we argue that changes in higher-order statistics such as in the covariance structure of data have to be manifest in the data before any conclusion about coordinated coding across voxels can be made.

In this study, Geodesic distance that reflects the underlying non-Euclidean geometry of covariance matrices [37] is used for measuring sMVR. This metric considers both rotation and scale differences between covariance matrices and outperforms other alternatives such as S-statistic which is the sum of orientation statistic and shape statistic [38] and Box’s M test [39]. S-statistic is very sensitive to outliers, because it works based on variance. M test is very sensitive to departures from normality, and requires data to be multivariate normally distributed.

For measuring MVR, Crossnobis distance that is a bias-free, reliable, and interpretable measure of dissimilarity was used [2, 40]. Alternatively, one may use ross-validated multivariate analysis of variance (CV-MANOVA) [41] that is a powerful method for comparing multivariate sample means. Although both CV-MANOVA and Crossnobis distances take into account the covariance matrices, none of them track changes in the covariance matrix across two conditions. On the other hand, the mean differences play a more important role in these methods. The Geodesic distance on the other hand considers covariance matrices explicitly without considering mean differences to evaluate the role of covariance matrices in decoding of task conditions.

Notably, analysis of real fMRI data in two species failed to show a significant sMVR coding within regions that nevertheless had strong MVR effects or anywhere else in the brain. Importantly, theoretical predictions based on the forward hemodynamic models using simulations revealed inherent limitations for fMRI signals to contain detectable sMVR even at unrealistically high SNR for conventional fMRI studies. Our results in this study call for caution in interpreting significant MVPA results as representing anything beyond a collection of univariate effects. It remains to be seen whether the advent of high field MRI (7T and above) with higher spatial resolutions and use of more sophisticated analysis for extracting higher-order interactions between voxels can overcome the current limitations in finding ‘true’ modulation of multivoxel coordination by task conditions of interests.

## 8. Methods

## 9. Crossnobis distance

In this dissimilarity measure, the data is divided into two independent samples, and the Mahalanobis distance is computed in each split separately [28]. Consider *X*_*A*_ ∈ ℝ *N*×*V* (*X*_*B*_ ∈ ℝ *N*×*V*) represent *N* samples of condition *A* (*B*) with *V* Voxels.

The first parameters that need to be estimated are the means and the covariance matrices of conditions. The mean of condition *A* is estimated as 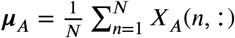 and the covariance matrix can be estimated as

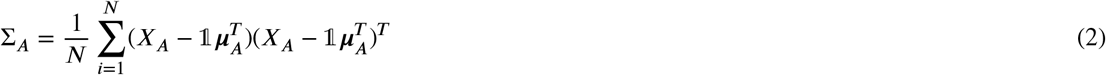

where ***µ***_*A*_ ∈ ℝ ^*V* ×1^, Σ_*A*_ ∈ ℝ ^*V* ×*V*^ and 1 = [1, …, 1] ∈ ℝ ^*N*×1^. Σ_*B*_ is computed in the same way.

In this cross-validated method, each matrix is divided into two independent sub-matrices *R* times (Fig S1A) and then in each *r*th partition (*r* = 1, …, *R*), four different Mahalanobis distances are computed as following

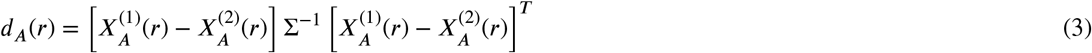

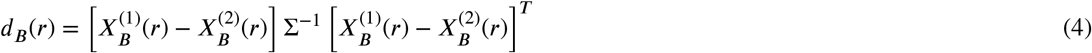

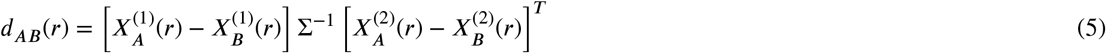

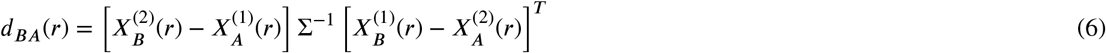

where 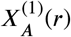 and 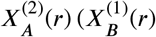and 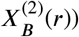are two independent sub-matrices for *X*_*A*_ (*X*_*B*_) in *r*th random partitioning. Note that equal covariance matrices for two conditions are assumed in this measure (Σ _*A*_ = Σ _*B*_ = Σ). In addition, if the covariance matrix of two conditions are not equal in reality, then the average of covariance matrix of *X*_*A*_ and *X*_*B*_ is assigned to Σ.

Thus, Crossnobis distance as MVR extraction method is defined as following

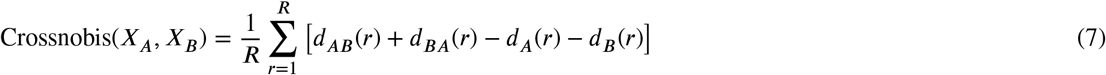

For real fMRI datasets, Crossnobis(*X*_*A*_, *X*_*B*_) is divided to the number of voxels in that searchlight to eliminate the effect of searchlight size. Note that this measure with applying cross-validation technique removes biases originated from noise [41], therefore, Crossnobis distance is more reliable than simple distance measures [28]. On the other hand, since this distance does not have an upper limit unlike the accuracy criterion [13], it is a better measure to represent the ease of decoding for the two conditions (Fig. 1B).

Crossnobis distance can quantify the impact of noise correlation in a voxel set on the decoded information about conditions (Fig S1B). It should be noted that in the literature, noise correlation illustrates the degree to which the variability of responses (in repeated presentations of the same condition) is shared between pairs of voxels [25, 42]. As a result, Crossnobis distance takes into account noise correlations that have an important impact on the accuracy of any decoding method [43].

## 10. Geodesic distance

Geodesic distance (Geodesic distance) is a geometry-aware approach in comparing covariance matrix structures and uses the positive semi-definiteness of covariance matrices to represent the underlying non-Euclidean geometry of them (Fig S2A-B). This distance assumes that covariance matrices lie on a non-linear space and finds their shortest distance along this manifold. As a result Geodesic distance is a sensitive and of the most accuarte methods to examine changes in the covariance matrices [29]. For two covariance matrices Σ_*A*_ and Σ_*B*_, their Geodesic distance can be quantified as

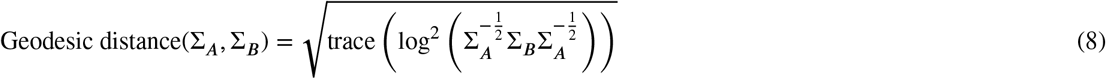

Since this measure is not symmetric between conditions A and B, we have used the average of Geodesic distance(Σ_*A*_, Σ_*B*_) and Geodesic distance(Σ_*B*_, Σ_*A*_) instead.

At the end, Geodesic distance(Σ_*A*_, Σ_*B*_) is divided to the number of voxels in that searchlight to eliminate the effect of searchlight size. Geodesic distance is applied in various fMRI analyses especially in comparing functional connectivity matrices [37, 44] and comparing representational similarity matrices in the brain and models [45].

Fig S2B shows Geodesic distance on the nonlinear manifold of positive semi-definite matrices. Note that, unlike the Euclidean distance, Geodesic distance constraints the distance of path to be within the given manifold which contains only valid covariance matrices (symmetric positive semi-definite matrices) [37].

To reduce the effect of small sample size on estimating the correct covariance matrices, the covariance matrices were estimated via more efficient methods such as [46, 47] and then the computed matrices were compared with Geodesic distance. The method in [46] considers the maximum likelihood (ML) estimation for the covariance matrix besides applying constraints on the positive semi-definiteness, as well as enforcing condition number upper-bounds. Furthermore, the method in [47] is a distribution-free estimator that leads to a closed-form covariance estimate. While the reported results are not done with these robust methods, using these methods did not change any of the main findings (i.e. lack of sMVR in fMRI data or simulations).

## 11. Permutation test

Estimating the statistical significance of detected differences between two conditions needs to be approached with care due to the high dimensionality of the data and the relatively small number of samples [48]. Permutation tests are one type of non-parametric test and their essential concept is relatively intuitive. If there is no experimental effect, then the labeling of samples by the corresponding condition is arbitrary, because the same data would have arisen regardless of the condition. Consequently, the null hypothesis would be the labeling is arbitrary and the significance of a main statistic (obtained from original data) can then be assessed by comparison with the distribution of values obtained when the labels are permuted.

If *N* is the number of random relabelings in the permutation test, then the P-value is:

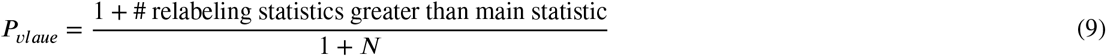

In this study, for each decoding task in both MVR and sMVR extraction methods, *N* = 999 and the reported results have *uncorrected* Pvalues less than 0.01.

For the group-level analysis of MVR and sMVR for the human dataset (Figs. 3B and 4B), average Crossnobis and Geodesic distance for each voxel was averaged across all 17 subjects and this observed average was compared with the distribution of averages obtained from the *N* = 999 relabelings for each subject to calculate group-level P-value for each voxel.

Minimum cluster size of 30 voxels was used for reported results for both human and monkey datasets for both single subject and group-level analyses.

## 12. Simulated fMRI data

To investigate the presence of MVR and sMVR in real fMRI data, first blood oxygenation level-dependent (BOLD) signal is simulated using forward hemodynamic model [49]. The forward hemodynamic model predicts the BOLD signal one would expect to measure with given neural activity and it explains the procedure that generates BOLD time-series from the underlying causes. This model applies a detailed biophysical model of the generation of the

BOLD signal based on the Balloon model [50]. The pathway from neural activity to BOLD signal can be divided into (1) interactions of neuronal activity and neurovascular coupling (flow induction), (2) blood vessel dynamics (Balloon model) [51], and (3) these parts combine to generate the observed nonlinear BOLD signal [52]. A brief explanation about this model is depicted in Fig 7A. Furthermore, the effect of spatial integration across neural populations due to large voxel sizes is also considered. Therefore, before applying DCM forward model, some random sets of neurons are selected from the neuron pool, and the average of the responses of these neurons is then applied to the hemodynamic forward model.

It this study, the MATLAB routines of freely available *SPM* toolbox [53] (SPM 12) is used to simulate BOLD signal based on DCM hemodynamic model. The main functions that are used from SPM toolbox are *spm_fx_fmri*.*m* and *spm_gx_fmri*.*m* with default hemodynamic parameters (repetition time= 2, rate of signal decay (,*c*) = 0.64, stiffness of blood vessels (*α*) = 0.32, autoregulation (*γ*) = 0.32, hemodynamic transit time (*τ*) = 2.00, resting oxygen extraction (*E*_0_) = 0.4, time to echo (*T*_*eco*_) = 0.04, ratio of intra-and extravascular signal (*є*) = 0.5, resting venous volume (*V*_0_) = 4).

Note that neuronal activations are not the only contributors to BOLD signals. The sensitivity of fMRI data analysis methods in detecting effects is susceptible to the relative levels of signal and noise in the BOLD signal. The sources of noise in fMRI data are related to instabilities in the recording system, subject head motion, and physiological fluctuations [54, 55]. Therefore, noise with a predefined signal-to-noise ratio (SNR) is added to the synthesized BOLD signal.

The input neural data to synthesize BOLD signal was taken from a simultaneous extracellular recording of neuronal populations in visual areas V1 and V2 [56, 57] a previously published and publically available dataset (http://crcns.org). In this study, the V2 recordings are used that were performed using tetrodes and include five recording sessions in three animals. The presented stimuli in this dataset are sets of oriented gratings of 8 different orientations (22.5 interval) in a pseudorandom sequence ^1^.

The real V2 data for every pair of different grating orientations (*0*_*A*_ and *0*_*B*_) were entered into the model and corresponding simulated BOLD signal is generated. Besides, the function *spm_rand_power_law*.*m* is applied to generate random variables with a power law spectral density, and the noise is added to simulated BOLD signal with different SNRs such that 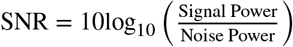.

## 13. fMRI data preprocessing

The preprocessing of two datasets (monkey and human) was performed in AFNI software packages [58, 59]. The preprocessing steps are as following:

1. warping anatomy to standard space (*auto_warp*.*py*)
2. aligning each dataset to base volume to correct head motion (*3dvolreg*)
3. aligning datasets to standardized anatomy (*3dNwarpApply*)
4. slice time correction (*3dTshift*)
5. removing spikes (*3dDespike*)

For sMVR analysis, first the GLM analysis using *3dDeconvolve* was carried out to conduct the beta coefficients using model time series and nuisance regressors [30]. The model time series consisted of two regressors ([right, left hemifield] for monkeys in left/right experiment, [face, object] for monkeys in the face/object experiment, and [face, scene] for human dataset) that were 1 during the condition’s blocks and 0 otherwise and were convolved with a MION hemodynamics. Seventeen nuisance regressors were motion and their first derivatives (12 parameters), blinks, and eye position (horizontal, vertical, and interaction).Before regression, all nuisance time series except the ones related to the motion were convolved with MION hemodynamics.

After preprocessing and GLM analysis, the first three samples of each condition’s blocks (preprocessed fMRI signal and GLM residuals) were discarded, and then the blocks of each task condition were concatenated to make one matrix for each condition. Then Crossnobis distance from preprocessed BOLD signal and Geodesic distance from GLM residuals were computed. Note that Geodesic distance is computed for each run separately and then averaged across runs. Also, a searchlight-based analysis using CoSMoMVPA toolbox [60] (with *V* = 15 voxels in each searchlight) is performed to obtain a brain wide map of activations.

## 14. Sources of data

The human fMRI dataset is available in the Princeton Neuroscience Institute data repository at http://dataspace.princeton.edu/jspui/ [Accessed: 2020-11-25]

## 15. Author Contributions

MP and AG conceived of the problem and the required analysis. MP performed the data analysis and simulations in discussions with AG. Both authors wrote the paper.

## Footnotes

1 This dataset is available at https://portal.nersc.gov/project/crcns/download/v1v2-1 [Accessed: 2020-06-20].

