## Supplemental Figures for "Coordinated multivoxel coding beyond univariate effects is not likely to be observable in fMRI data"

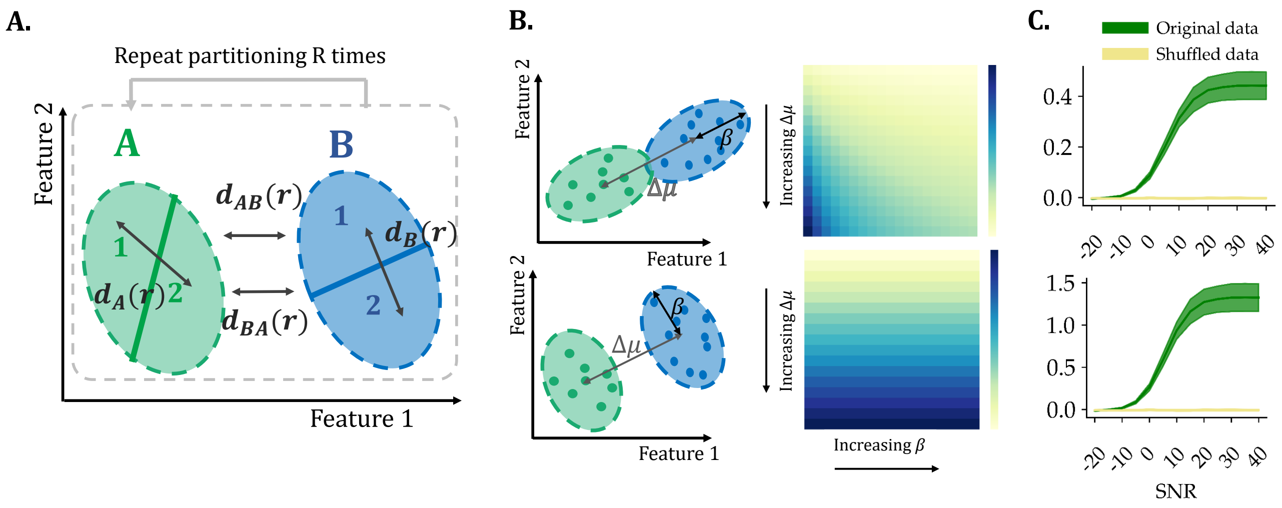


Fig. 1: (A) MVP extraction method using Crossnobis distance: the samples in each condition split into two independent partitions and four different Mahalanobis distances are computed. (B) Effect of changing $\Delta\mu$ and β on Crossnobis distance for correlated effect in two conditions (the eigenvector corresponding to the largest eigenvalue β is in the same direction as $\Delta\mu$) and for anti-correlated effect in two conditions (β is perpendicular to the $\Delta\mu$). Upper figures show that Crossnobis distance is sensitive to both $\Delta\mu$ and β when the mean response of two conditions are correlated. As it can be seen, with increasing $\Delta\mu$, Crossnobis distance increases, but with increasing β, Crossnobis distance decreases because the samples of the two classes collapse and become difficult to separate. On the other hand, if the mean response of two conditions is anti-correlated with eigenvector corresponding largest eigenvalue, the Crossnobis distance is not sensitive to parameter β (Fig. 1B lower) because increasing the variance of the samples in the direction perpendicular to the line between means does not cause the samples of the two classes to be confused. (C) Crossnobis distance for simulated data versus different SNRs in dB for correlated (upper) and anti-correlated (lower) effect.

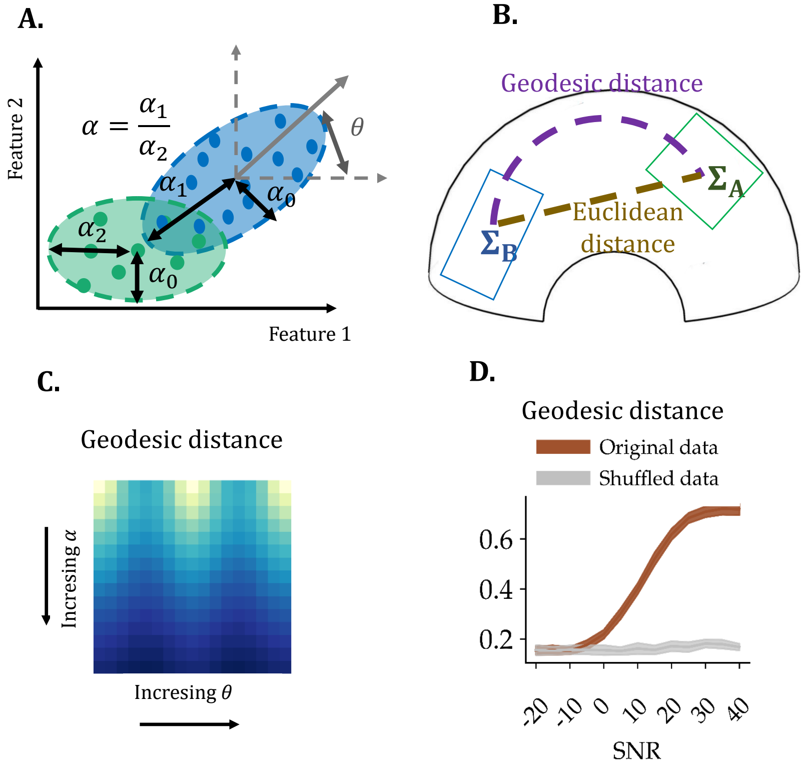


Fig. 2: (A) Important parameters in simulated data including θ (difference in orientation between covariance matrices) and α (difference in shape) (B) Geodesic distance and Euclidean distance on the reimanian manifold. (C) Effect of changing θ and α on Geodesic distance. The Geodesic distance is sensitive to changes in both α and θ. (D) Geodesic distance when Gaussian noise with different SNRs in dB is added to data for $\theta \neq 0$ and $\alpha \neq 0$. As it can be seen, the more noisy the data, the more likely it is that there will be outliers, and outliers will lead to drastic changes in data distribution (in terms of orientation and shape).

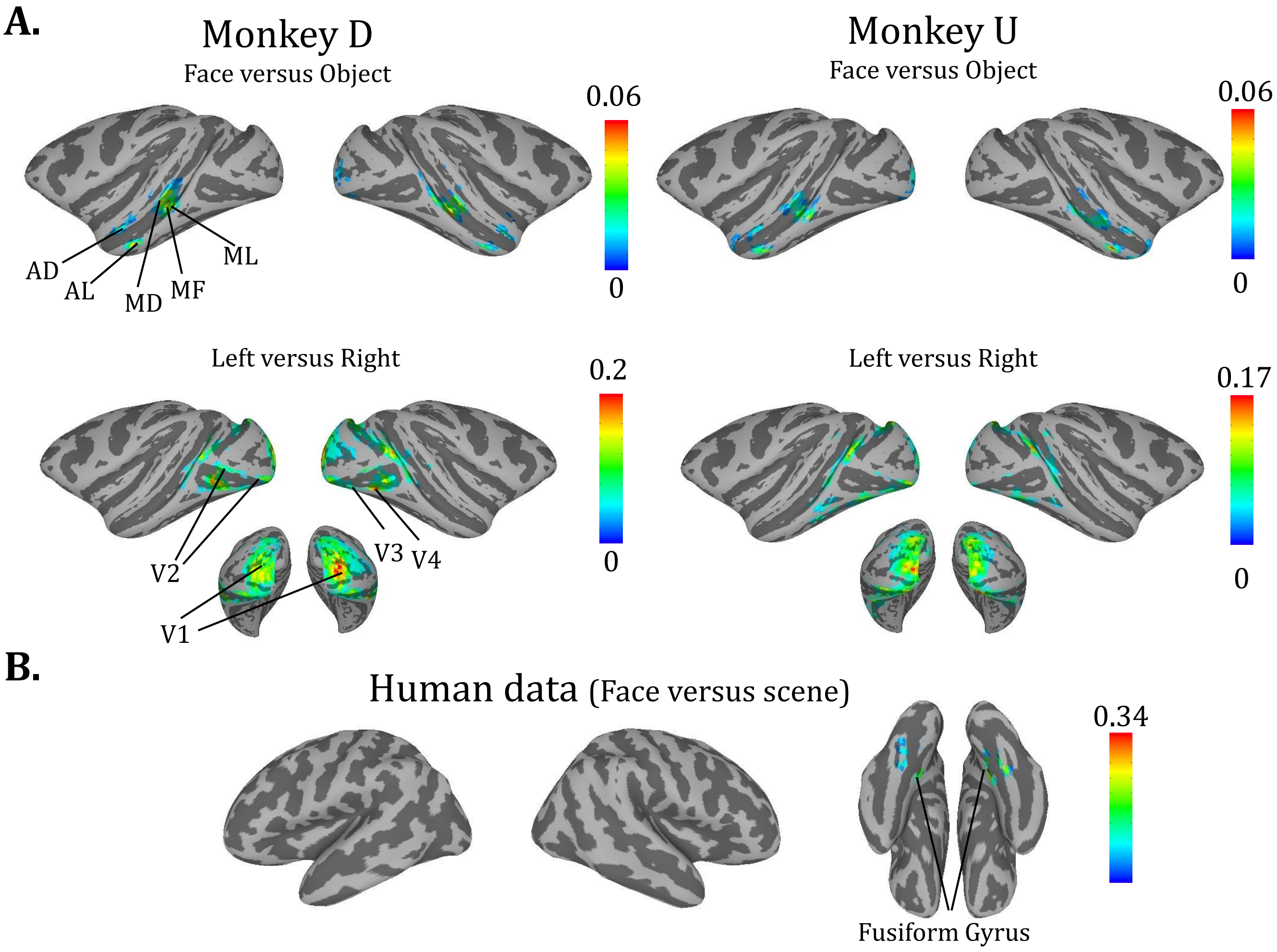


Fig. 3: Object discrimination using general linear model across cortical areas. (A) Face versus Object and Left versus Right beta coefficients in areas with significant discrimination for monkeys D and U. (B) The average of Face versus Scene beta coefficients in areas with significant discrimination for 17 subjects. For all plots $P < 0.01$, with minimum cluster size 30.

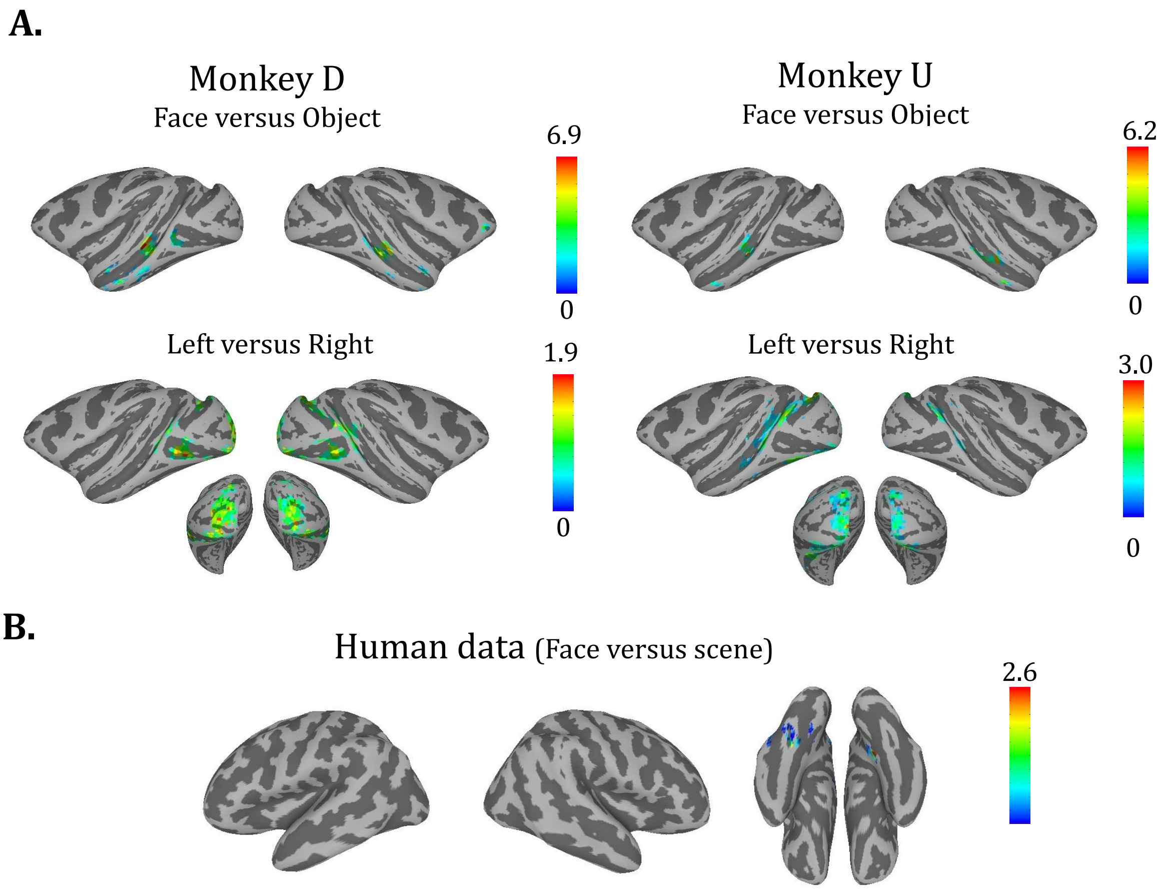


Fig. 4: UVR for face/object and left/right hemifield across cortical areas (A) Crossnobis distance (with 1 voxels in each searchlight) for two monkeys for face/object and left/right discrimination tasks. The highlighted regions show significant cluster of voxels ($P_{value} = 0.01$, cluster corrected, cluster size 30) . (B) The average of Crossnobis distance of 17 subjects for face/scene discrimination task. The highlighted regions are significant cluster of voxels ($P_{value} < 0.01$ with minimum cluster size 30).

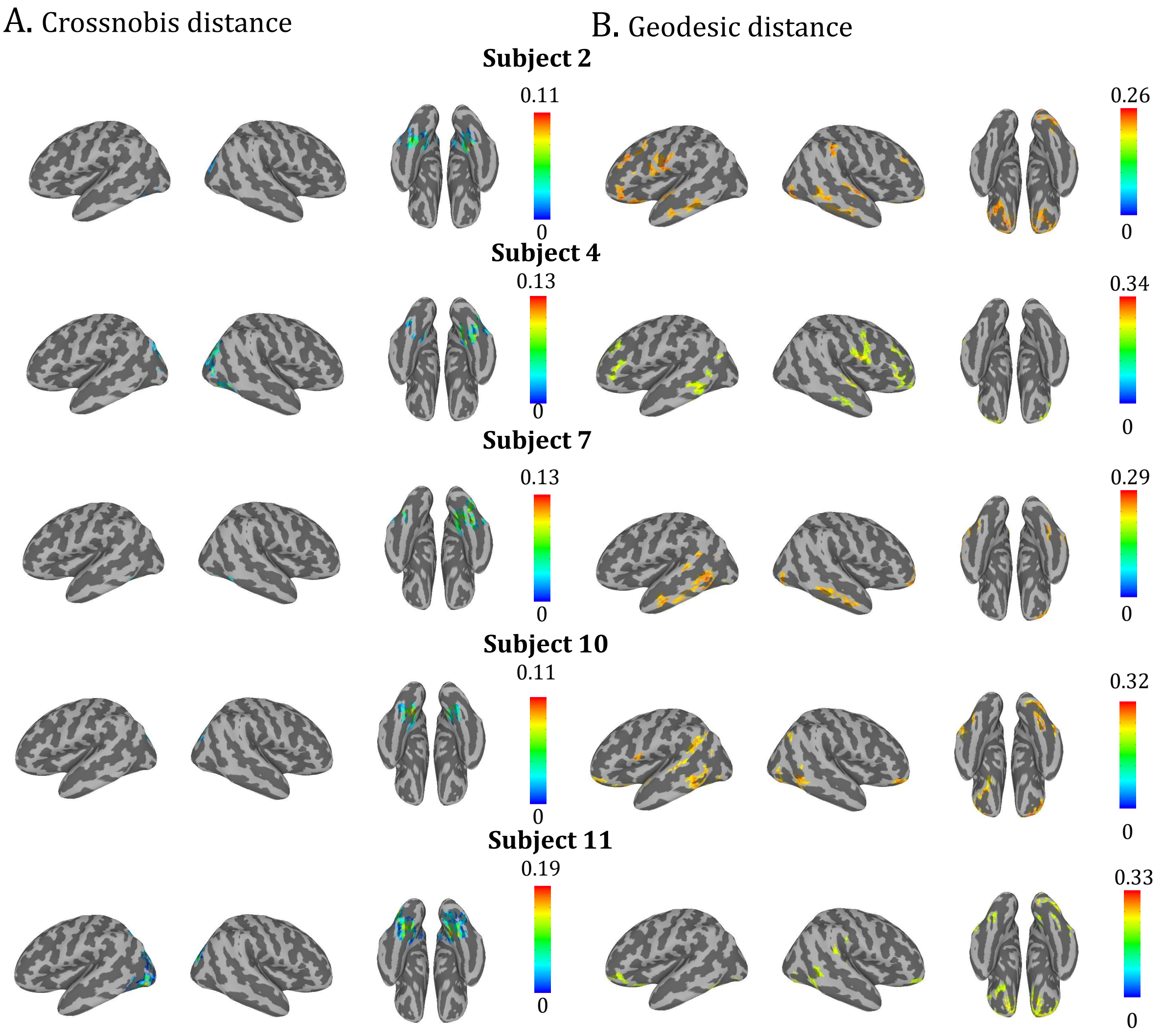


Fig. 5: sMVR and MVR for face/scene across cortical areas of five subjects in human dataset (A) Crossnobis distance with 15 voxels in each searchlight, $P_{value} < 0.01$, minimum cluster size 30. (B) Geodesic distance The highlighted regions show top 20% regions with highest sMVR and cluster correcting (cluster size=30). Note that none of clusters in (B) were significant ($P_{value} > 0.01$).

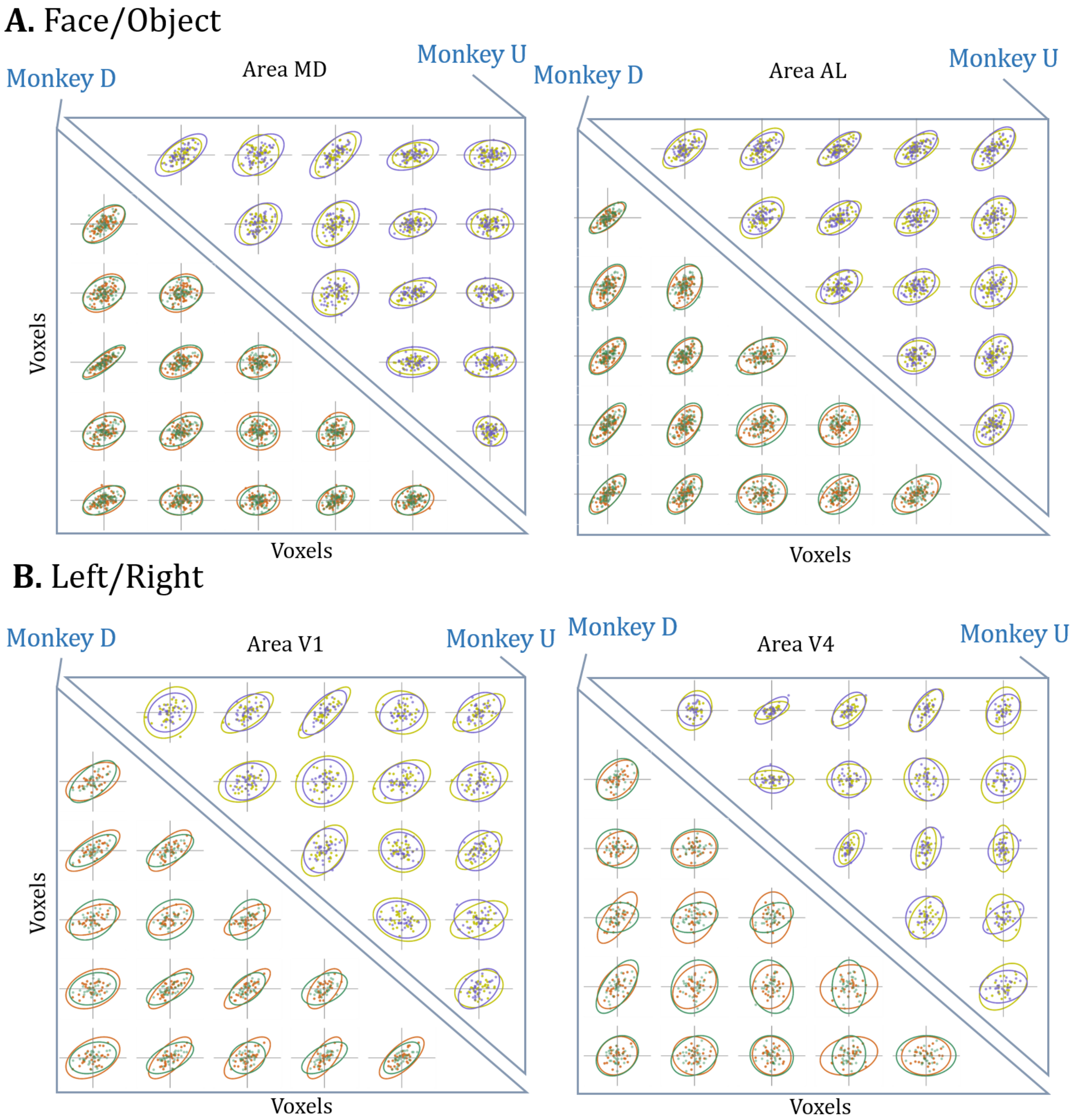


Fig. 6: Visualization of changes in covariance of voxels across conditions with removing mean of each task condition. (A) Joint-distribution of pairwise voxel activations in monkey D (lower triangle) and monkey D (upper triangle) in face vs object blocks for two different searchlight in area MD (left) and area AL (right) with significant Crossnobis distance and non significant Geodesic distance. (B) same format as A but for left vs right blocks in area V1 (left) and area V4 (right).

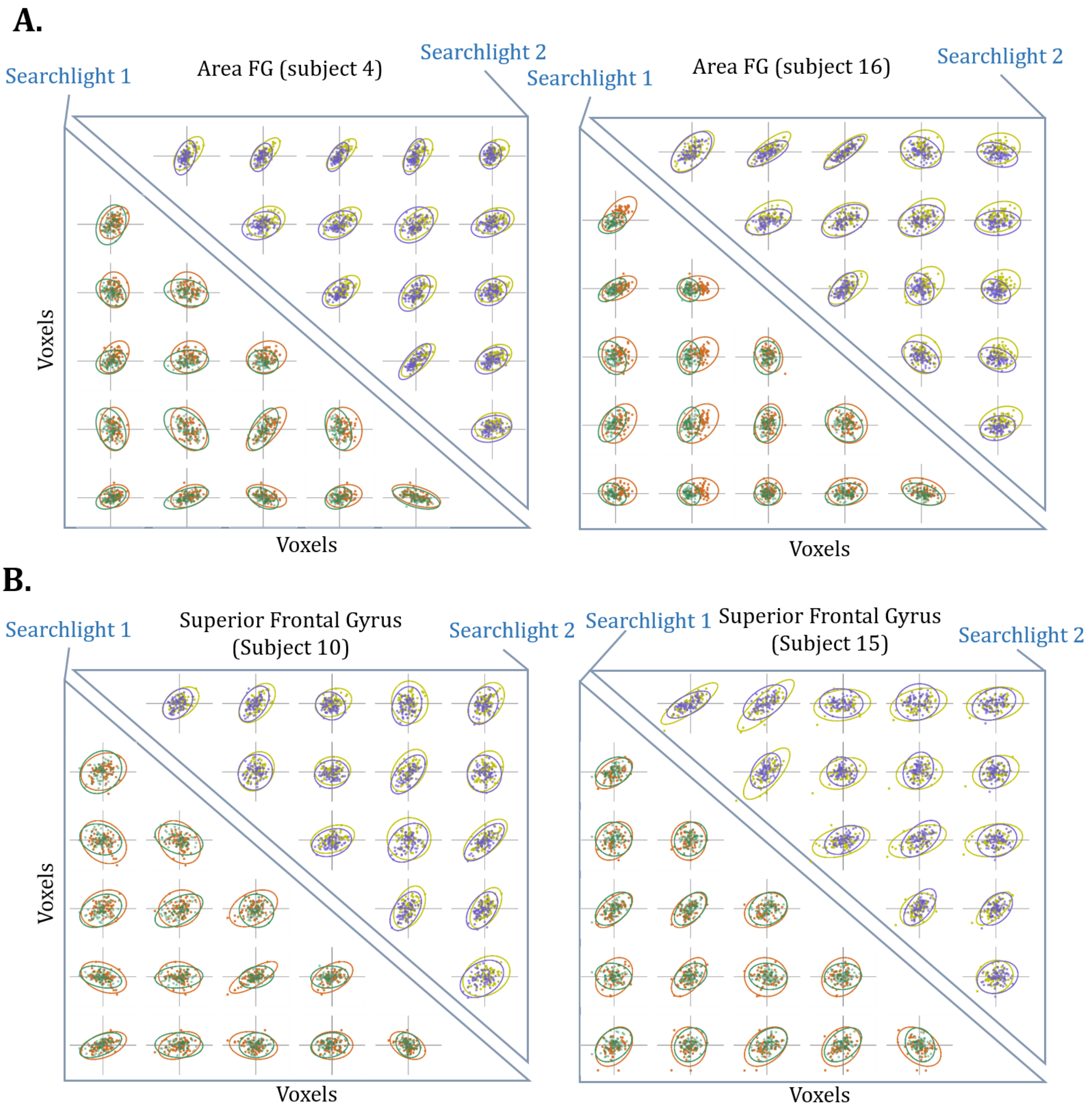


Fig. 7: Visualization of changes in covariance of voxels across conditions for human dataset. (A) Joint-distribution of pairwise voxel activations in two different searchlights (lower triangle and upper triangle) in face vs scene blocks in area V1 for subject 4 (left) and subject 16 (right). In these examples, MVR is significant but sMVR is not significant. (B) same format as A but neither MVR and sMVR are not significant. Different conditions are plotted with different colors.

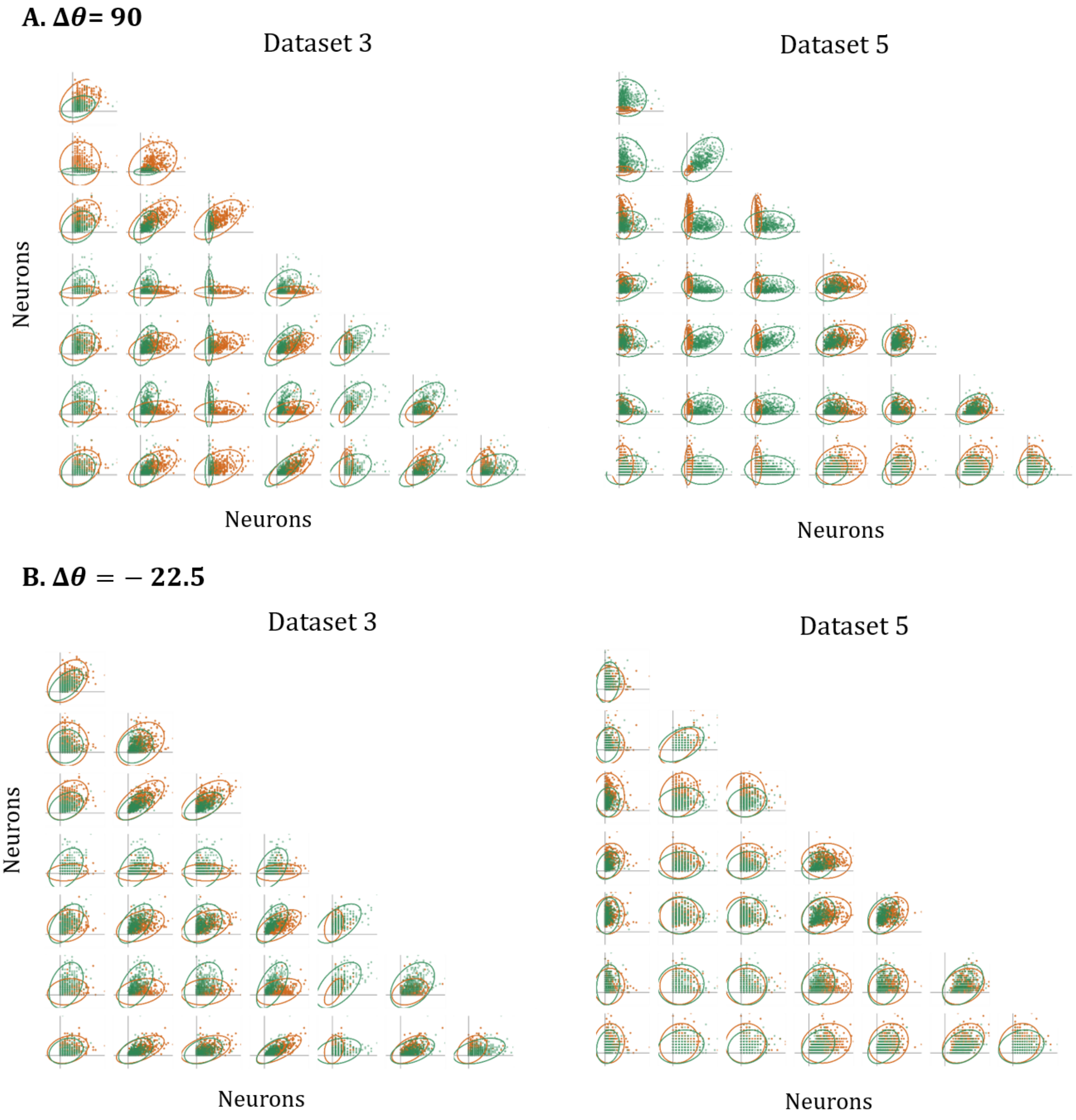


Fig. 8: Visualization of changes in covariance of activity of neurons across conditions. (A) Two examples of distribution of samples for $\Delta\theta = 90$ for datasets 3 (left) and 5 (right). Different colors corresponds to two different $\theta_A$ and $\theta_B$ (B) same format as A but for for $\Delta\theta = -22.5$ for datasets 3 (left) and 5 (right). The lower and upper triangle are the distribution of samples recorded in V2 and V1 area, respectively.

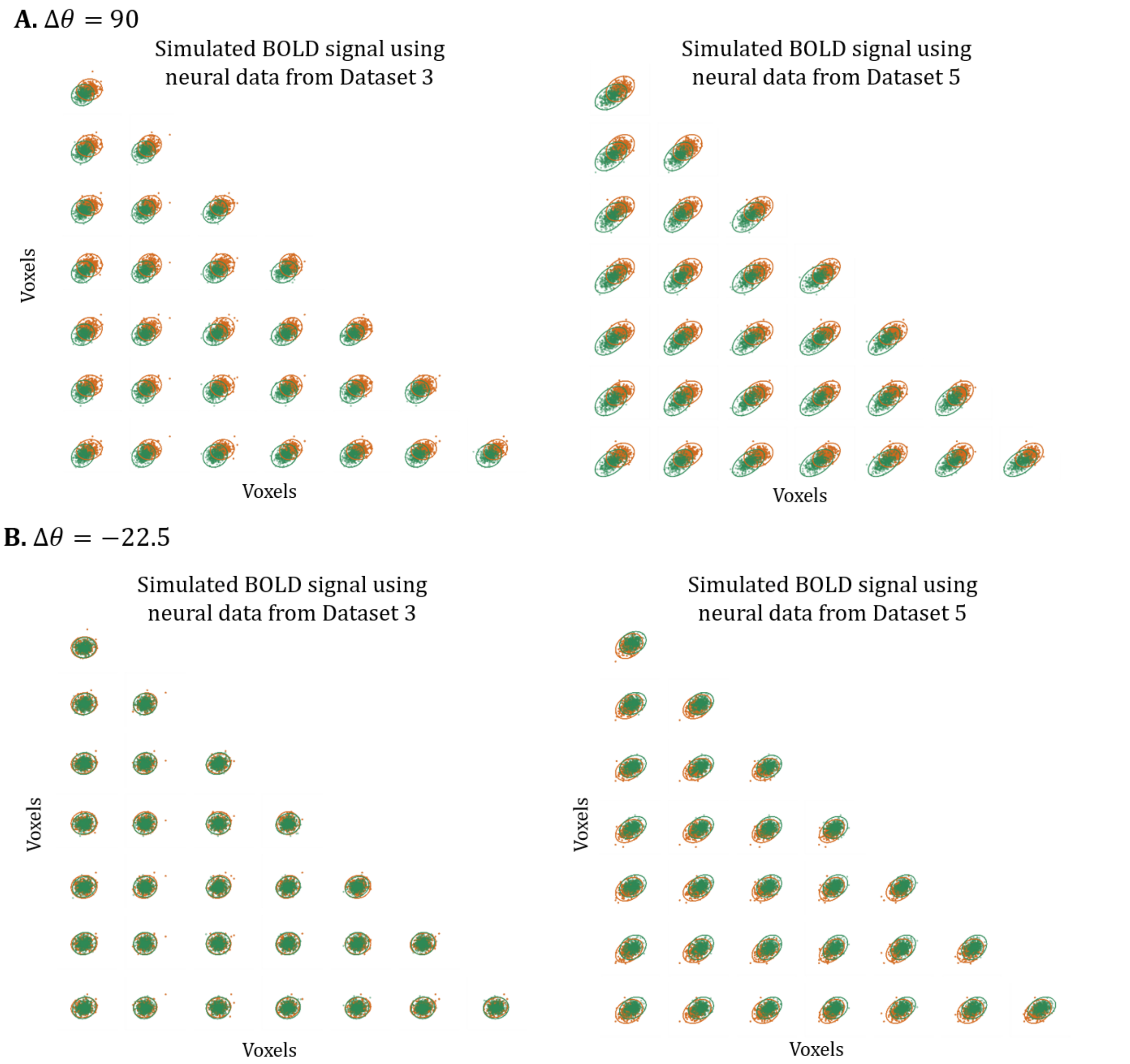


Fig. 9: Visualization of changes in covariance of simulated BOLD sample. (A) Two examples of distribution of simulated fMRI data using real neural data from datasets 3 and 5 for $\Delta\theta = 90$. Each condition's samples are plotted with specific color. (B)same format as A but for $\Delta\theta = -22.5$. In all of plots SNR = 10dB.

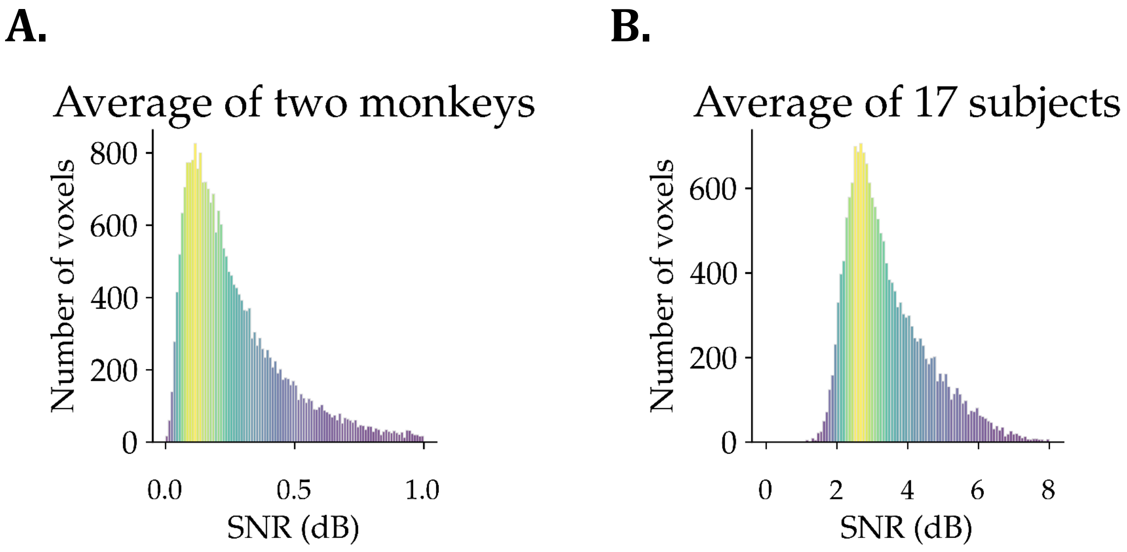


Fig. 10: Histogram of SNR values for (A) the average of two monkeys and (B) the average of 17 human subjects.
